# A self-initiated cue-reward learning procedure for neural recording in rodents

**DOI:** 10.1101/2020.02.24.963447

**Authors:** Ingrid Reverte, Stephen Volz, Fahd H. Alhazmi, Mihwa Kang, Keith Kaufman, Sue Chan, Claudia Jou, Mihaela D. Iordanova, Guillem R. Esber

**Author notes:** Fondazione Santa Lucia, Via del Fosso di Fiorano 64, 00143, Rome, Italy (Ingrid Reverte). Corresponding author: Guillem R. Esber.

## Abstract

**Background:** Single-unit recording in Pavlovian conditioning tasks requires the use of within-subject designs as well as sampling a considerable number of trials per trial type and session, which increases the total trial count. Pavlovian conditioning, on the other hand, requires a long average intertrial interval (ITI) relative to cue duration for cue-specific learning to occur. These requirements combined can make the session duration unfeasibly long.

**New Method:** To circumvent this issue, we developed a self-initiated variant of the Pavlovian magazine-approach procedure in rodents. Unlike the standard procedure, where the animals passively receive the trials, the self-initiated procedure grants animals agency to self-administer and self-pace trials from a predetermined, pseudorandomized list. Critically, whereas in the standard procedure the typical ITI is in the order of minutes, our procedure uses a much shorter ITI (10 s).

**Results:** Despite such a short ITI, discrimination learning in the self-initiated procedure is comparable to that observed in the standard procedure with a typical ITI, and superior to that observed in the standard procedure with an equally short ITI.

**Comparison with Existing Method(s):** The self-initiated procedure permits delivering 100 trials in a ∼1-h session, almost doubling the number of trials safely attainable over that period with the standard procedure.

**Conclusions:** The self-initiated procedure enhances the collection of neural correlates of cue-reward learning while producing good discrimination performance. Other advantages for neural recording studies include ensuring that at the start of each trial the animal is engaged, attentive and in the same location within the conditioning chamber.

## 1. Introduction

Progress in behavioral neuroscience rests on the foundation of well-controlled behavioral designs capable of isolating the cognitive process of interest and yielding replicable results (Krakauer et al., 2017). However, adapting traditional behavioral procedures to meet the requirements of neuroscience techniques may demand some ingenuity. One class of challenge stems from the fact that experimental parameters favorable to the cognitive process under investigation may conflict with those that best suit the neuroscience technique at hand. This conflict becomes apparent, for instance, when investigating the neural correlates of cue-reward learning with neural recording techniques such as in-vivo electrophysiological recording and calcium imaging. Here, we introduce a self-initiated, self-paced conditioning procedure for rodents specifically designed to enhance the acquisition of neural data in such scenarios.

The mechanisms of cue-reward learning have been dissected by learning theorists using Pavlovian conditioning procedures (Mackintosh, 1974; Kehoe & Macrae, 2002), which have led and continue to lead to highly influential findings in neuroscience (e.g., Hawkins et al. 1983; Kim et al., 1998; Maren, 2001; Schultz & Dickinson, 2000; Waelti et al., 2001; Holland, 1997). In the rat, one such procedure is the conditioned magazine approach (e.g., Boakes, 1977; Harris et al., 2013), in which animals receive presentations of certain cues or conditioned stimuli (CSs; typically, visual or auditory) followed when appropriate by the delivery of a reward or an unconditioned stimulus (US; e.g., sucrose solution). Reward expectancies are typically quantified by measuring the total number of head-entries or the cumulative percentage of time spent in the reward magazine during the CS before the US is delivered (Gottlieb, 2005). A discrimination is said to emerge as the rat responds more in the presence of rewarded than unrewarded cues. Critically, as in other Pavlovian procedures (Prokasy & Ebel, 1964; Salafia et al., 1973; Terrace et al., 1975; Domjan, 1980; Gibbon & Balsam, 1981; Yin et al., 1994; Barella, 1999), better performance is observed with spaced rather than massed trial presentations; that is, when the intertrial interval (ITI) is sufficiently long relative to the duration of the cues (i.e., the trial-spacing effect; Lattal, 1999; Holland, 2000).

Although scheduling a long ITI benefits learning, it can be problematic in neural recording studies. To illustrate why, consider the results of a series of bootstrap analyses conducted on in-vivo electrophysiological data recorded in the rat orbitofrontal cortex (Fig. 1 and Supplemental Materials). The data were collected during the CS epochs of a well-trained visual discrimination using the novel reward-learning procedure introduced here. The top panels show that the number of neurons that significantly discriminate between rewarded and unrewarded trials steadily increases as more trials are sampled. The bottom panels bolster this point by showing that the statistical power observed for each neuron in the same analysis is also a monotonically-increasing function of the number of trials sampled. The story told by this figure will be familiar to many in-vivo electrophysiologists: single-unit recording requires the presentation of a sizeable number of trials per trial type in order to average out the trial-to-trial variability inherent to neural data.

**Figure 1.**
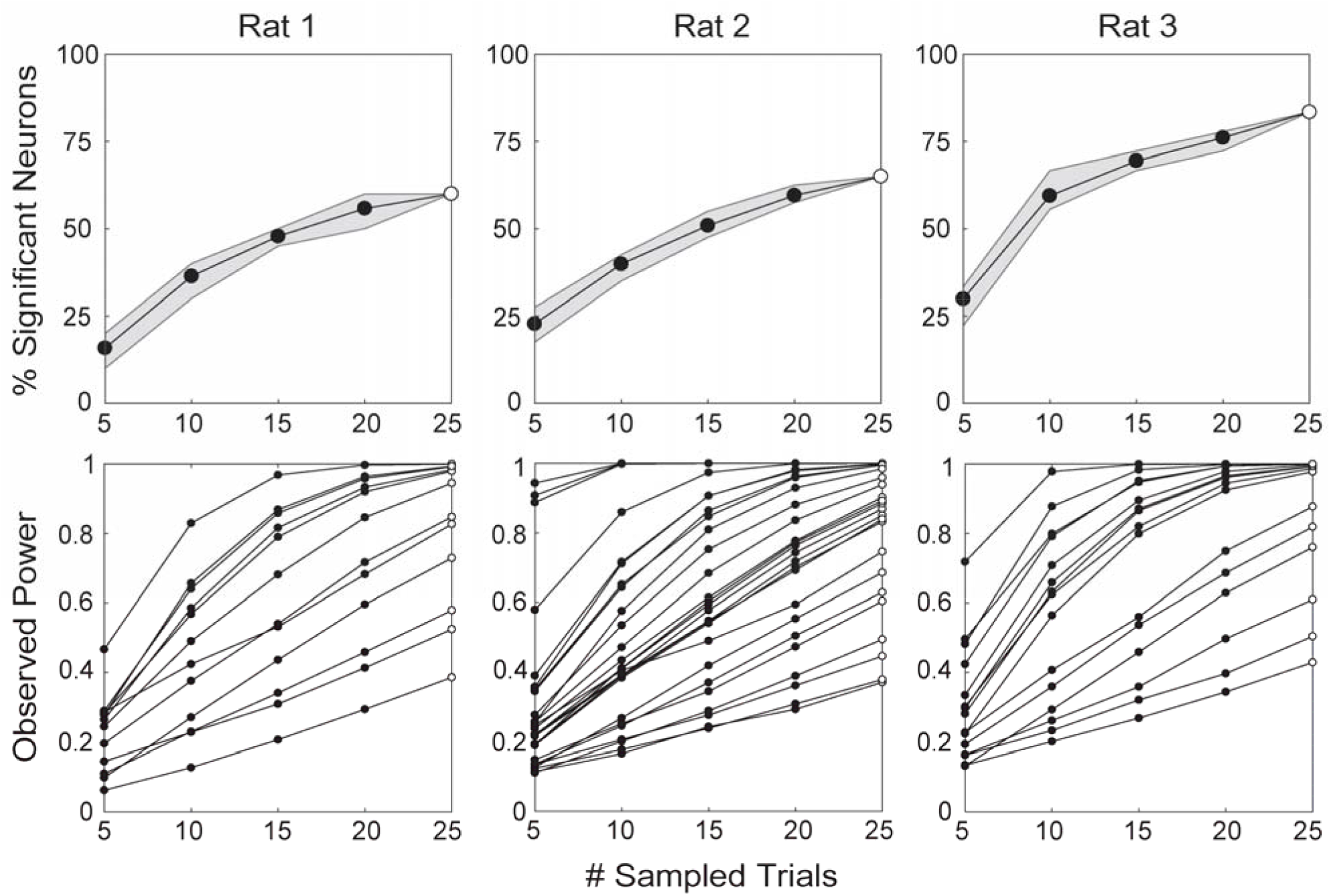
Results of a series of bootstrap analyses demonstrating the importance of large trial counts for investigating the neural correlates of predictive learning (see Supplemental Materials). All analyses were conducted using spike rates during CS period of neurons recorded in the orbitofrontal cortex of three rats (columns) on the final session of discrimination training of the form V1+, V2-, V1V2-, where V1 and V2 represent two 10-s visual cues, while the “+” and “–“ symbols represent reinforcement and non-reinforcement, respectively. Each trial type was presented 25 times in a session, adding up to a total of 75 trials. The top panels show the percentage of neurons that significantly discriminated between reinforced and non-reinforced cues as a function of the number of trials sampled, as identified by a one-way ANOVA (p<0.05). Shaded areas represent the 75% and 25% quartiles of the bootstrap iterations. The bottom panels depict the mean observed statistical power in the same ANOVA for each neuron recorded, also plotted as a function of the number of trials sampled. Open circles represent the actual results when all 25 trials presented were included.

This requirement, when combined with that of a long ITI in Pavlovian procedures, will produce lengthy neural recording sessions—often unfeasibly so once the experimenter ventures beyond basic discrimination designs. To compound the issue, neural recording studies demand the use of within-subject designs in order to compare neural responses between experimental and control cues, further contributing to elevating the total trial count in a session. This makes examining the neural bases of discrimination, categorization and rule learning difficult for the in-vivo electrophysiologist working with rodents. Such scenarios involve more complex experimental designs, leaving the experimenter with a hard choice between shortening the ITI, which can jeopardize learning, and reducing the number of trials at the peril of insufficiently sampling neural activity.

To circumvent this choice, we developed a variant of the Pavlovian conditioned magazine-approach procedure we have dubbed the *self-initiated conditioned magazine approach* (SICMA) procedure. Unlike the standard procedure, where the rat passively receives the trials, in SICMA it falls upon the animal to initiate each trial by performing a separate response upon receiving a cue signaling trial availability. Because the ITI is only 10 s on average, SICMA permits packing 100 trials in a ∼1 h session, almost doubling the number of trials safely attainable in that time with the standard Pavlovian procedure. Crucially, despite such a short ITI, our results show that performance in SICMA is comparable to that observed in the standard procedure (Experiment 1), and superior to that observed in a yoked Pavlovian group (Experiment 2). In addition, we provide evidence that magazine-approach responses to cues trained with the SICMA procedure readily transfer when the cues are presented in a standard Pavlovian fashion (Experiment 3). Thus, SICMA affords the in-vivo electrophysiologist an opportunity to efficiently examine the neural underpinnings of cue-reward learning using complex discrimination designs.

## 2. Experiment 1: comparison of SICMA with the standard Pavlovian magazine-approach procedure

The goal of this experiment was to compare a group of rats trained with the SICMA procedure (labeled SICMA) with another one trained with the standard Pavlovian magazine-approach method (labeled Pav) in their ability to solve two discriminations involving visual and auditory cues. The experimental parameters used in the Pav group (e.g., ITI and CS durations) were known from prior unpublished work from our laboratory to produce good discrimination performance.

### 2.1. Materials and Methods

#### 2.1.1. Animals

All animal care and experimental procedures were conducted according to the National Institutes of Health’s *Guide for the Care and the Use of Laboratory Animals*, and approved by the Brooklyn College Institutional Animal Care and Use Committee (Protocol #303). Subjects were 32 experimentally-naïve, adult Long-Evans rats (16 males and 16 females) bred at Brooklyn College from commercially available populations (Charles River laboratories). At the start of the experiment, all rats were approximately 90 (+/-7) days old and their weights ranged between 244 and 271 g for females and between 317 and 340 g for males. They were housed individually in standard clear-plastic tubs (10.5 in. × 19 in. × 8 in) with woodchip bedding in a colony room on a 14:10 light/dark schedule. Behavioral sessions were conducted between 3-6 hours after the onset of the light phase of the cycle. Throughout training, food was provided *ad libitum* but water access was restricted to 1 h/day immediately after each experimental session.

#### 2.1.2. Apparatus

Behavioral training was conducted in eight standard conditioning chambers (Med Associates Inc., St. Albans, VT, USA) measuring 32 cm in length, 25 cm in width and 33 cm in height, and comprising a stainless-steel grid floor, a Perspex front door, back wall, and ceiling, and modular left and right walls. Each chamber was enclosed in a ventilated sound-attenuating cubicle (74 cm x 45 cm x 60 cm) that provided a background noise level of ∼50 dB. A schematic depiction of the interior of the chambers is shown in Figure S1 (Supplemental Materials, Section S2.1). All reported locations of stimulus and response apparatus were measured from the grid floor of the conditioning chamber to the lowest point or edge of the apparatus. The left wall of the chamber housed two white jewel lamps 2.5 cm in diameter (28V DC, 100 mA) located 9.3 cm above the grid floor on the left and right panels, as well as a speaker (7 cm x 8.2 cm) located 20.6 cm above the grid floor on the right panel and connected to a dedicated tone generator capable of delivering a 12-kHz, 70-dB tone. The right wall housed a third white jewel lamp (28V DC, 100 mA) 2.5 cm in diameter, located 17.2 cm above the grid floor on the center panel, as well as a speaker located 24.8 cm above the grid floor on the left panel and connected to a dedicated tone generator capable of delivering a 70-dB white noise. The right wall also housed a circular noseport 2.6 cm in diameter located on the center panel 4.6 cm above the grid floor, equipped with a yellow LED light and an infrared sensor for detecting nose entries. This noseport was flanked by a recessed liquid reward magazine (aperture: 5.1 cm x 15.2 cm) located on the right panel 1.6 cm above the grid floor. This magazine was equipped with an infrared sensor for detecting head entries, and connected to a liquid dipper that could deliver a 0.04 cc droplet of a 10% sucrose solution. The chambers remained dark throughout the experimental session except during presentations of the visual stimuli. In the same room was a computer running Med PC IV software (Med Associates Inc., St. Albans, VT, USA) on Windows OS which controlled and automatically recorded all experimental events via a Fader Control Interface.

#### 2.1.3. Procedure

##### 2.1.3.1. Magazine training and shaping

Animals were first randomly assigned to the SICMA and Pav groups (16 rats in each, gender balanced). Each session began with a 2-min acclimation period in the conditioning chambers. Rats were initially magazine-trained in a 1-h session to retrieve up to 60 deliveries of a 10% sucrose reward at the dipper magazine. For the first 10 trials, the reward was made available for 30 s every 30 s; for the second 20 trials, it was available for 20 s every 40 s; and finally, for the last 30 trials, it was available for 10 s every 50 s.

Rats in the SICMA group then went on to receive five additional shaping sessions. On the first of these sessions, the noseport light was turned on for a maximum of 20 s, during which a nose poke immediately resulted in the termination of the noseport light and the onset of the sucrose reward, which remained available for 10 s. Trials were separated by a 10 s variable ITI (range: 5-15 s). Over the following four shaping sessions, we introduced and progressively increased a delay (2, 4, 6, and 8 s) between the rat’s response at the port and reward delivery, during which the noseport light would flash at a 1-Hz frequency (on for 0.5 s, off for 0.5 s). Concurrently, reward availability was progressively shortened (8, 6, 4, and 3 s).

##### 2.1.3.2. Trial structure

Fig. 2 depicts the basic trial structure in the SICMA procedure. As during shaping, rats in the SICMA group were still required to self-initiate trials in this phase by responding at the lit-up noseport during the 20-s periods of trial availability. Failure to respond resulted in the noseport light coming off and the trial being repeated after a short ITI averaging 10 s and ranging 5-15 s. In contrast, performing a nose-poke response immediately terminated the noseport light and triggered the onset of one of four possible 10-s CSs. Reinforced trials culminated in 3 s of access to the sucrose reward, followed by a short ITI (average 10 s; range: 5-15 s). In contrast, rats in the Pav group received the 10-s CSs in the standard Pavlovian conditioning manner (i.e., noncontingent on any response), followed, whenever reinforced, by the same reward used in the SICMA group. The ITI in the Pav group was 60 s on average (range: 40-80 s).

**Figure 2.**
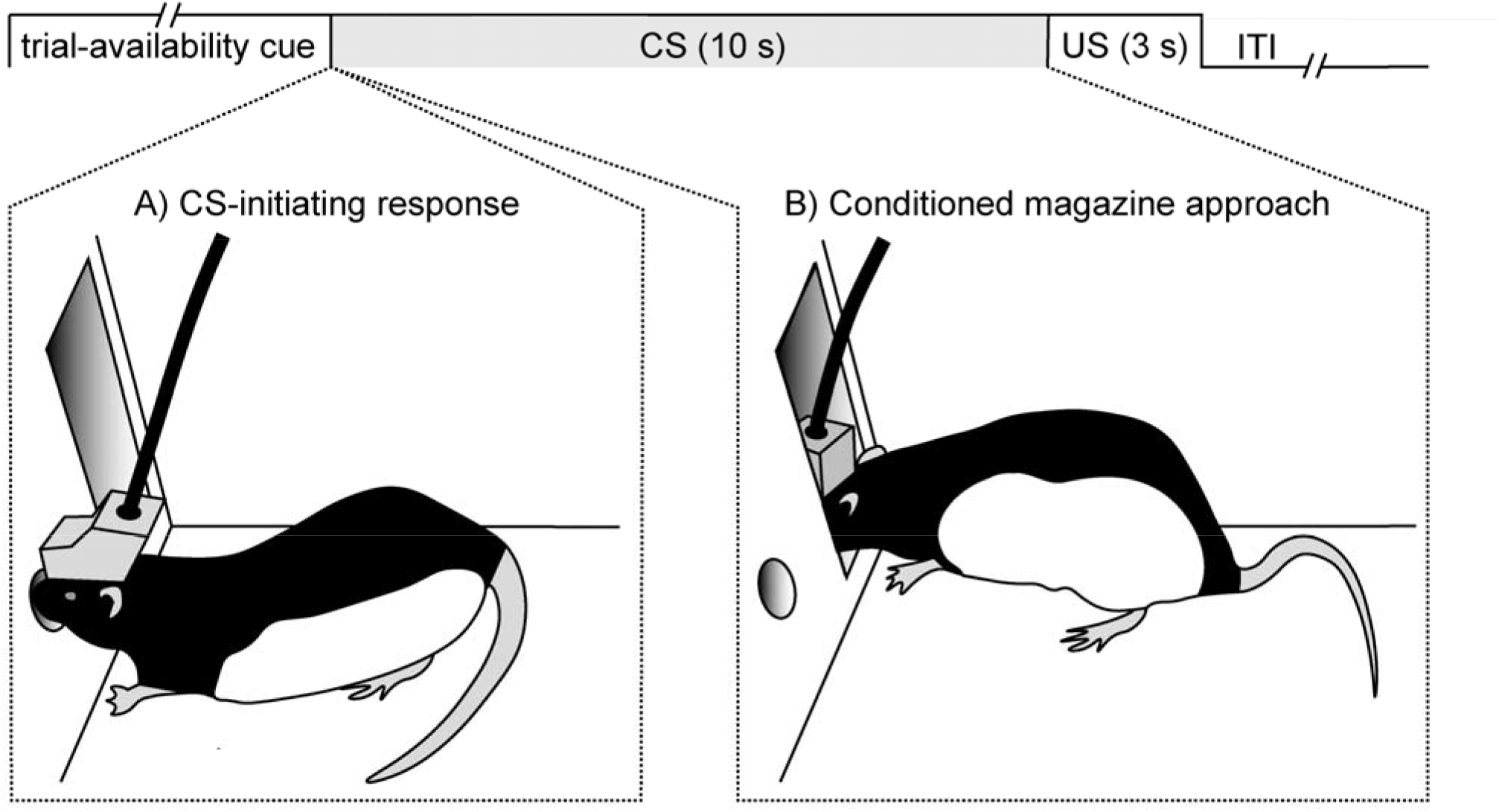
Trial schematic of the SICMA procedure. A light cue inside the noseport signals trial availability for a maximum of 20 s, during which the animal can respond at the noseport (panel A) to turn on one of several possible CSs. During the 10-s CS, the animal may perform anticipatory approach responses in the reward magazine (panel B)—just as in the standard magazine-approach procedure. On reinforced trials, a reward (US) is delivered at the end of the CS, followed by an average intertrial interval of 10 s.

##### 2.1.3.3. Discrimination training

Although any discrimination can be imbedded in SICMA, the experiments reported here involved two discriminations, one involving two visual CSs (V1 & V2, counterbalanced) and the other two auditory ones (A1 & A2, counterbalanced). A table containing the details of the stimulus counterbalancing can be found Section S3 of the Supplemental Materials (Table S1). One visual CS was constructed by flashing the two jewel lamps on the left wall alternately at a 2-Hz frequency (on for 0.25 s, off for 0.25 s). The second visual CS was provided by the steady illumination of the white jewel lamp located on the right wall. The two auditory CSs were provided by a 12-kHz, 70-dB tone played from the speaker on the left wall and a 70-dB white noise played from the speaker on the right wall. The probability of reinforcement varied across the CSs, with V1 and V2 reinforced on 100% and 0% of trials, respectively, and A1 and A2 reinforced pseudorandomly on 75% and 25% of trials, respectively. In the SICMA group, each session ended when the rat completed 96 trials or else it timed out at 90 min. Rats in the Pav group received a total of 64 trials per session. Although this may seem an unfair comparison from the viewpoint of trial-centered theories of predictive learning (e.g., Rescorla & Wagner, 1972; Wagner, 1981; Stout & Miller, 2007), evidence indicates that the number of trials in a session has no measurable effect on the rate of acquisition when assessed—as in the present case—in between-subject designs (Gottlieb, 2008).

##### 2.1.3.4. Statistical analysis

For this and the remaining experiments, we used the percentage of time each rat spent in the reward magazine during the cues, a widely used measure of conditioned responding (e.g., Kaye & Pearce, 1984; Hunt & Campbell, 1997; Holland, 1999; Gottlieb, 2005). We chose this dependent variable above the other conventional measure—the rate of head entries per minute—because we have observed that in SICMA-trained rats it provides a more sensitive index of discrimination learning. This can be readily appreciated in Figure S2 (Section S2.2 of Supplemental Materials), which depicts the count of rats in the SICMA and Pav groups across the last two sessions of Experiment 1 as a function of the mean number of head entries during cues V1 (left panel) and A1 (right panel)—the cues with the highest reinforcement probability within either sensory modality.

The figure shows rather different response distributions for each cue between the groups. Specifically, the distributions are more positively skewed in the SICMA (skewness: V1 = 1.7, A1 = 1.7; kurtosis: V1 = 1.8, A1 = 2.3) than the Pav group (skewness: V1 = 1.2, A1 = 0.8; kurtosis: V1 = 1.3, A1 = −0.2), with the majority of SICMA observations consisting of a single response. Indeed, the median response rate in the SICMA group for both cues was 1, whereas that in the Pav group was 2.7. A Mann-Whitney test confirmed that rats in the SICMA group made fewer head entries than those in the Pav group both during V1 (U = 222, p<0.0001) and A1 (U = 220, p<0.0001). Such a low response variability in SICMA-trained rats discourages the use of rate of head entries as a dependent variable in SICMA studies, and confines any conclusions drawn from group comparisons here to percent responding. In any case, it is worth noting that we (unpublished) and others (e.g., Takahashi et al., 2013) have found that, likely due to the physical restraint imposed by the tether, electrode-implanted animals also express discrimination learning more clearly in percent responding than rate of head entries in the Pavlovian magazine approach procedure.

For the purpose of statistical analyses, the data from each subject was first averaged across trials in a session, and further collapsed into average responding in two-session blocks. Analyses of the cues A1/A2 and V1/V2 were conducted separately, as these two subsets of cues differ in both modality and probability of reward, making comparisons across cue pairs uninformative. Results were analyzed using a mixed-model linear analysis ANOVA, and Bonferroni-corrected simple-effects analysis to decompose significant interactions when present. All calculations were conducted in JAMOVI (Gallucci, 2017; The Jamovi Project, 2019).

### 2.2. Results and Discussion

Overall, SICMA rats completed all trials on 94% of the sessions. The mean session duration was 73.1 min (SD = 25.1) in the SICMA group and 77.5 min in the Pav group. The effective mean ITI in the SICMA group (10-s ITI + latency to nose poke after trial-availability cue onset + 30-s no-initiation trials) was 21 s (SD = 11.8 s).

#### 2.2.1. Comparison of the temporal dynamics of magazine approach between the groups

Due to task requirements, SICMA rats started each trial with their nose in the noseport and thus had a constant distance (∼8 cm) to travel to enter the adjacent reward magazine (Fig. 2). In contrast, the distance between rats in the Pav group and the magazine at the start of each trial could vary. To investigate the impact of such differences in starting location at cue onset on the topography of conditioned responding, we calculated the second-by-second percentage of time spent in the magazine during the visual (Fig. 3, top left panel) and auditory (Fig. 3, top right panel) cues in the final 2-session block of training. Overall, rats in the SICMA group responded more to both the visual (F_Grp_(1,30)=7.91, p=0.009) and auditory (F_Grp_ (1,30)=7.43, p=0.011) stimuli, compared to rats in the Pav group. This difference between groups emerged across the duration of the cues (visual: F_Grp*Sec_(9,570)=182.57, p<0.001; auditory: F_Grp*Sec_ (9,570) = 15.96, p<0.001). With regard to the visual stimuli, only V1(100%) produced differential levels of responding between the two groups (F_Grp*Sec*CS_(9,570)=2.37, p=0.012). In the first second following CS onset the groups did not significantly differ in their response to V1(100%) (F(1,82.4)=0.42, p≈1) or V2(0%) (F(1,82.4)=0.90, p≈1), but for all subsequent seconds the SICMA group showed significantly higher levels of responding to V1(100%) than the Pav group (F(1,82.4)=14.40-25.08, Ps<0.02. Although this suggests a higher response ceiling in the SICMA group, it is worth noting that the groups did not differ in their ability to withhold responding in the presence of V2(0%) (F(1,82.4)=0.002-1.04, Ps≈1 in all seconds beyond the first). Thus, the higher response ceiling for V1(100%) in the SICMA group does not appear to result from an indiscriminate elevation of baseline responding in these animals.

**Figure 3.**
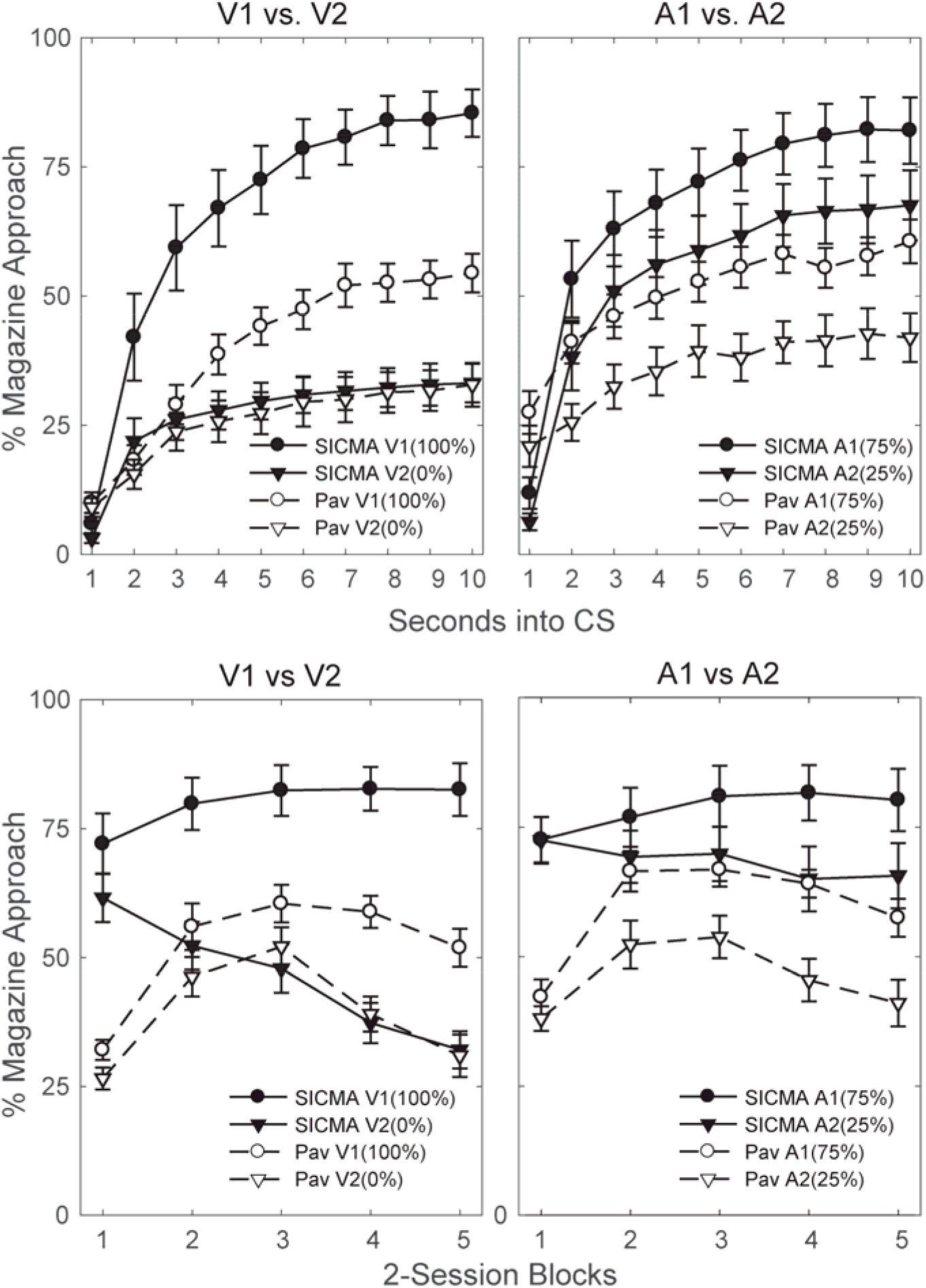
Comparison of conditioned magazine-approach performance in a visual (left panels) and auditory (right panels) discrimination between the SICMA and Pavlovian groups. The top panels show the time course of responding to the CSs in the final 2-session block of training, expressed as the mean percentage of time the rats spent in the magazine in each of the 10 s of cue presentation. The bottom panels show the mean percentage of time the rats spent in the magazine during the 10-s CSs across the five 2-session blocks of discrimination training. Error bars represent the standard error of the mean (SEM).

As for the auditory discrimination, no overall between-group differences in responding to stimuli A1(75%) and A2(25%) were detected in the first 5 s of cue period (F(1,43.3)=3.21-7.91; p>0.08). Notably, responding was numerically greater in the Pav than the SICMA group in the very first second, presumably indicating that some Pav rats may have been near or even inside the reward magazine at the time of cue onset— a physical impossibility for SICMA animals. Greater responding to these cues in the SICMA relative to the Pav group did reach significance in the sixth second, and stayed significant for the remainder of the auditory cues period (F(1,43.3)=10.22-13.41, p<0.03). Thus, this result suggests that the SICMA procedure might encourage greater responding to partially reinforced cues, at least from the auditory modality.

#### 2.2.2. Comparison of discrimination learning between the groups

To compare discrimination learning between the groups, we analyzed magazine activity during the cues across the five two-session blocks of training (Fig. 3, bottom panels). Following Holland (1977), we focused our analysis on the last 5 s of CS period, where a more stable readout of magazine activity can be obtained (Fig. 3, top panels). The results, shown in the bottom panels of Fig. 3, confirmed that rats across the two groups solved both the visual (F_CS_(1,270)=322.67, p<0.001) and auditory discriminations (F_CS_(1,270)=75.005 p<0.001).

Unsurprisingly, the solution of the visual discrimination emerged in both group as training progressed (F_Blk*CS_(4,270)=14.87; p<0.001; Fig. 3, bottom left panel). More importantly, this discrimination was solved more readily by the SICMA than the Pav group. (F_Grp*Blk*CS_(4,270)=2.89, p=0.023). Simple effects analyses revealed that the SICMA group showed significant evidence of discrimination learning between the visual cues from session block 2 onwards (F(1,270)=45.18-151.67, p<0.015). In contrast, the Pavlovian group only showed significant evidence of discrimination learning starting on session blocks 4 and 5 (F(1,270)=23.41-26.15, p<0.015). Additionally, there was a significant difference between the groups in overall level of responding on the first block of training (F(1,53.6)=53.86, p<0.015).

As expected, the auditory discrimination (Fig. 3, bottom right panel) similarly emerged over the course of training in both groups (F_Blk*CS_(4,270)=3.910, p=0.004). Simple effects analysis showed that, combined, both groups responded significantly more to A1(75%) than A2(25%) from the second session block onwards (F(1,270)=13.132-34.313, p<0.01). Due to the lack of a significant three-factor interaction, it is safe to interpret this finding as indicating that both groups solved the auditory discrimination by the second block of training and did not significantly differ from each other in their ability to discriminate the cues. There was, however, a significant difference between the groups in baseline levels of responding F_Grp_(1,30)=15.229, p < 0.001), which changed over the course of training (F_Blk*Grp_(4,270)=6.374, *p*<0.001). Simple effects analysis of this interaction showed that the groups significantly differed in their overall level of responding in blocks 1,4 and 5 (F(1,47.1)=29.73, p<0.01; F(1,47.1)=9.72, p=0.03 and F(1,47.1)=15.86, *Ps*<0.01, respectively), but not in blocks 2 or 3 (F(1,47.1)=5.30-6.46, *Ps*>0.14). This baseline difference aside, the results of Experiment 1 show that, despite the short ITI, rats trained with the SICMA procedure showed no worse (and if anything, better) discrimination performance than rats trained with the standard Pavlovian magazine approach procedure.

## 3. Experiment 2 – Comparison of SICMA and yoked Pavlovian magazine-approach groups

This experiment aimed to provide a more direct comparison between the SICMA procedure and the Pavlovian magazine-approach method by imposing identical training conditions except for the requirement self-initiation. To this end, a yoked procedure was used in which animals in the Pavlovian group (labeled Yoked) received the exact same sequence of experimental events and, critically, at the same time, as their self-initiating counterparts in the SICMA group, ensuring an equal number of equally spaced trials.

### 3.1. Materials and Methods

#### 3.1.1. Animals & Apparatus

Eight male and eight female adult Long-Evans rats bred at Brooklyn College from rats of Charles River descent were used (Charles River Laboratories). At the start of the experiment, all rats were approximately 90 (+/−7) days old and their weights ranged between 239 and 253 g for females and 301 and 334 g for males. They were kept under the same husbandry conditions as described in Experiment 1. Experimental sessions were conducted between 3-5 hours after the onset of the light phase of the cycle. The apparatus used was that described in Experiment 1.

#### 3.1.2. Procedure

Animals were randomly assigned to two groups, labeled SICMA and Yoked (8 rats per group, gender balanced). In the SICMA group, magazine training, shaping and discrimination training procedures were identical to those used in Experiment 1. Following magazine training, rats in the Yoked group were each paired with a master rat in the SICMA group. This ensured that each rat in the Yoked group received the same exact sequence of events and at the same time as it was being experienced by its master rat in the SICMA group. This included noseport light illumination at the start of each trial-availability period in the SICMA group. Thus, the only difference between the two groups was that the yoked rats had no behavioral control over trial initiation. The results were analyzed with the same statistical tests used in Experiment 1.

### 3.2. Results and Discussion

SICMA rats completed all trials on 96% of the sessions (idem, of course, in the yoked rats). The session duration in the groups was 53.8 min on average, with a SD of 11.5 min. The effective mean ITI in the SICMA group (10-s ITI + latency to nose poke after trial-availability cue onset + no-initiation trials) was 20.4 s (SD = 7.3 s).

#### 3.2.1. Comparison of the temporal dynamics of magazine approach between the groups

To examine potential differences in response topography due to between-group differences in the rats’ distance to the reward magazine at cue onset, once again we analyzed the temporal profile of magazine approach across the 10 s of CS presentation, focusing on the final two-session block of training (Fig. 4, top panels). Overall, rats showed changes in responding over the 10 s for both the visual (F_Sec_(9,266)=11.239, p<0.001) and auditory (F_Sec_(9,266)=8.117, p<0.001) discriminations. Interestingly, this effect of Second into stimulus presentation interacted with the Group factor for both visual (F_Grp*Sec_(9,266)=5.192, p<0.001) and auditory modalities (F_Grp*Sec_(9,266)=3.212, p=0.001). Indeed, the top panels of Fig. 4 show that rats in the SICMA group progressively increased responding after the first second of the better predictor in each discrimination, whereas rats in the Yoked group were consistent across its duration, an observation that was confirmed by simple effects analysis for both the visual (F_SICMA_(9,266)=15.07 p<0.004 and F_Yoked_(9,266)=1.36 p=0.816) and auditory modalities (F_SICMA_(9,266)=10.345, p<0.004 and F_Yoked_(9,266)=0.984, p≈1). Thus, this finding indicate a greater dynamic range of responding for SICMA than Pav subjects under the present training conditions (i.e., short ITI). The SICMA group responded less than the Yoked group in the first second of both discriminations, although this trend was not significant. Once again, this suggests that some of the Yoked animals were immediately adjacent to or inside the reward magazine at the time of cue onset, as would be expected given the short ITI.

**Figure 4.**
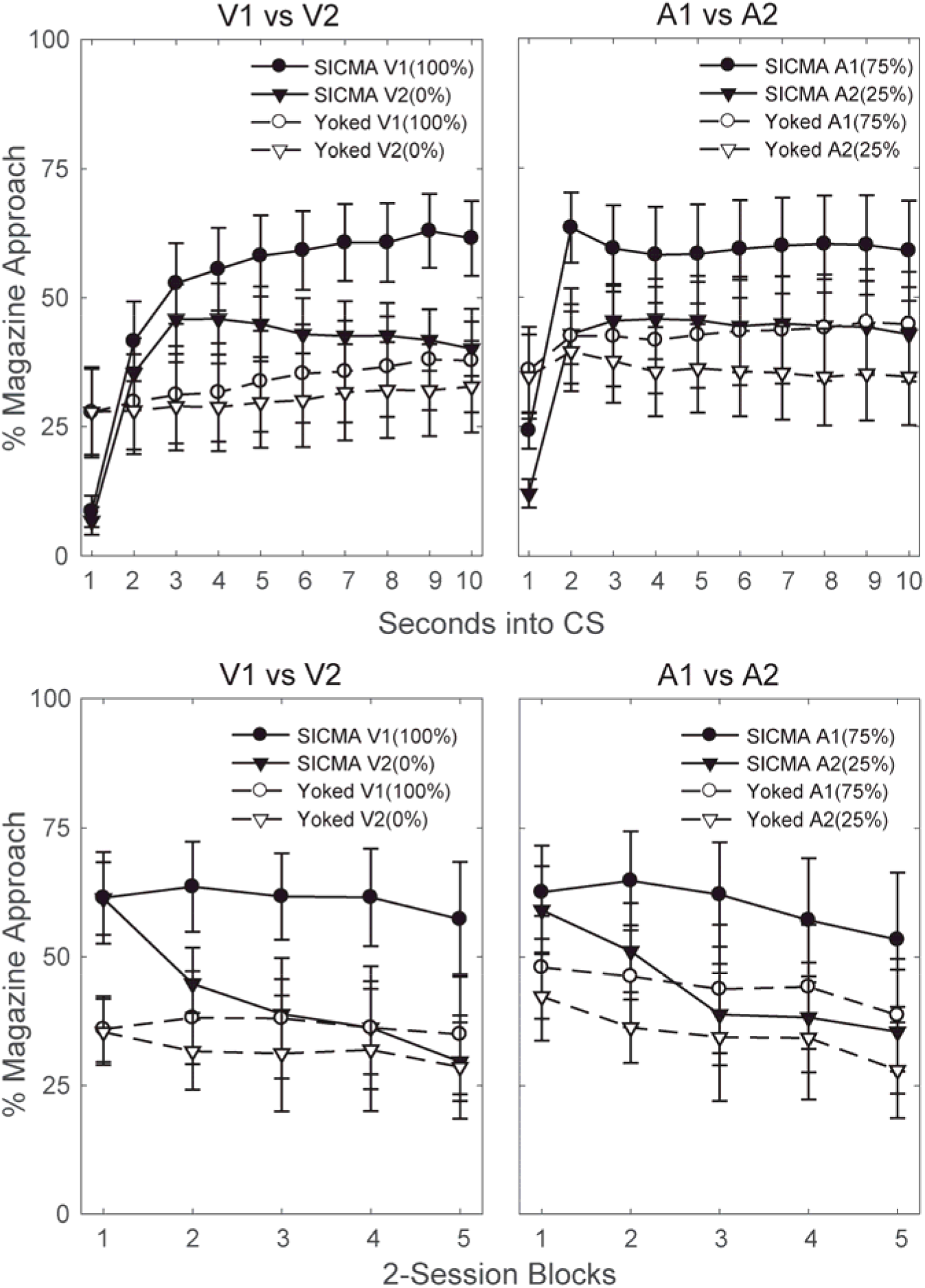
Comparison of conditioned magazine-approach responding in a visual (left panels) and auditory (right panels) discrimination between the SICMA and Yoked Pavlovian groups. The top panels depict the time course of responding to the CSs in the final 2-session block of training, expressed as the mean percentage of time the rats spent in the magazine in each of the 10 s of cue presentation. The bottom panels show the mean percentage of time the rats spent in the magazine during the last 5 s of CS period across the five 2-session blocks of discrimination training. Error bars represent the standard error of the mean (SEM).

#### 3.2.2. Comparison of discrimination learning between the groups

As in Experiment 1, to determine if and when the groups solved the two discriminations across training, we analyzed magazine activity across all five two-session blocks, focusing on the last 5 s period of CS presentation (Fig. 4, bottom panels). A main effect of Stimulus was significant in both modality discriminations (Visual: F_CS_(1,126)=26.697, p<0.001; Auditory: F_CS_(1,126) = 29.59, p<0.001), indicating that all rats considered together were able to discriminate between the cues as training progressed. Furthermore, a main effect of Session block was likewise significant (Visual: F_Blk_(4,126)=3.226; p=0.015; Auditory: F_Blk_(4,126)=8.4560, p<0.001), confirming that, as the bottom panels of Fig. 4 show, the discriminations were solved by withholding responding over the course of training to the less predictive CSs (V2 and A2) without increasing responding to the more predictive ones (V1 and A1).

Critically, as evident in the bottom panels of Fig. 4, the SICMA group showed better discrimination learning than the Yoked group, and this was true of the visual (F_Grp*CS_(1,126)=8.992, p=0.003) and auditory (F_Grp*CS_(1,126)=4.230, p=0.042) modalities. Indeed, the visual discrimination achieved statistical significance in the SICMA (F(1,126)=33.34, p<0.001), but not the Yoked group (F(1,126)=2.35; p=0.128). On the other hand, both the SICMA and Yoked groups solved the auditory discrimination to a significant degree (F_SICMA_(1,126)=28.10, p<0.002 and F_Yoked_(1,126)=5.72, p=0.036, respectively), although the SICMA animals solved this discrimination with a larger effect size (95% confidence interval of difference in percent responding: 4.662-10.22) than the Yoked rats did (0.580-6.13). Thus, discriminative performance in the Yoked group achieved significance in the case of the auditory, but not the visual discrimination, despite the latter being simpler in terms of the reward probabilities involved (100% vs 0% as opposed to 75% vs. 25% in the auditory case). This may simply reflect the superior perceptual discriminability of the auditory relative to the visual cues we used. Taken together, the results in the Yoked group confirm the deleterious effects of a short ITI in the conditioned magazine-approach preparation (e.g., Lattal, 1999; Holland, 2000)., and highlight the risk associated with shortening the ITI in neural recording studies using Pavlovian conditioning. Crucially, such deleterious effects were not observed in the SICMA group despite having an equally short ITI, the implications of which are considered in the General Discussion.

## 4. Experiment 3 – Does conditioned responding to self-initiated cues transfer when the cues are delivered in the standard Pavlovian fashion?

A notable difference between SICMA and the standard Pavlovian procedure is that SICMA requires shaping an instrumental nose-poke response at the noseport prior to the start of discrimination training. This raises the question of whether SICMA-trained rats come to treat the cues as Pavlovian CSs (i.e., cues that evoke Pavlovian conditioned approach responses) or rather as discriminative stimuli that inform the animal of when to complete an instrumental action sequence consisting of a nose poke followed by magazine approach. Although we would argue that neither associative structure would detract from the advantages of SICMA for neural recording, one particular scenario would render this procedure less useful. If during shaping rats acquire a noseport poke magazine approach action sequence, they could conceivably ignore reinforced CSs and learn only about cues that signal the omission of reinforcement. If this is the case, then reinforced cues trained with SICMA should evoke little magazine approach when delivered in a Pavlovian fashion (i.e., without self-initiation). In contrast, if reinforced cues trained with SICMA are attended to and learned about, such a transfer should be relatively seamless. Experiment 3 allows for the dissociation of these two possibilities.

### 4.1. Materials and Methods

#### 4.1.1. Animals & Apparatus

Four male and four female adult Long-Evans rats were used, bred at Brooklyn College from rats of Charles River descent. At the start of the experiment, all rats were approximately 90 (+/−7) days old and their weights ranged between 242 and 257 g for females and 311 and 345 g for males. Husbandry and apparatus details were identical to those reported in the previous experiments.

#### 4.1.2. Procedure

Magazine training, shaping and discrimination training procedures were identical to those used in the SICMA group of Experiment 1, except that animals received 20 sessions. The day after the last SICMA session, a single Pavlovian transfer session was conducted in which the rats were presented with the same discrimination. The procedural details in this test session were identical to those used in the Pav group of Experiment 2.

### 4.2. Results and Discussion

Trials were averaged into 2-trials blocks. We used a series of uncorrected within-subjects t-test to determine if performance in the Pavlovian transfer session was significantly different from that at final 2-trial block of SICMA training. We chose not to correct these t-test for multiple comparisons, as in this case we hypothesized that these conditions would not produce significant differences. As can be seen in Fig. 5, rats’ conditioned magazine activity to visual (top panel) and auditory (bottom panel) cues was virtually identical in the last 2-trial block of SICMA training and all-trial blocks of the Pavlovian transfer session. To ensure that these similarities were not due to rapid within-session acquisition, we focused our analysis on the first 2-trial block of the Pavlovian session. For the visual discrimination, t-tests found no significant differences in responding to V1 (t(7)=1.42 p=0.196) or V2 (t(7)=0.19 p=0.857), and these results were mirrored for the auditory cues (t(7)=2.08, p=0.075 for A1 and t(7)=-0.404, p=0.698 for A2). Thus, even under conditions favorable to detecting a difference (a series of uncorrected t-tests), the results confirm that the predictive significance of the cues was preserved when the cues were subsequently presented without self-initiation to animals that had never previously received Pavlovian training. This is inconsistent with the hypothesis that SICMA training discourages rats from attending to and learning about reinforced cues, at least when the latter are embedded in a discrimination.

**Figure 5.**
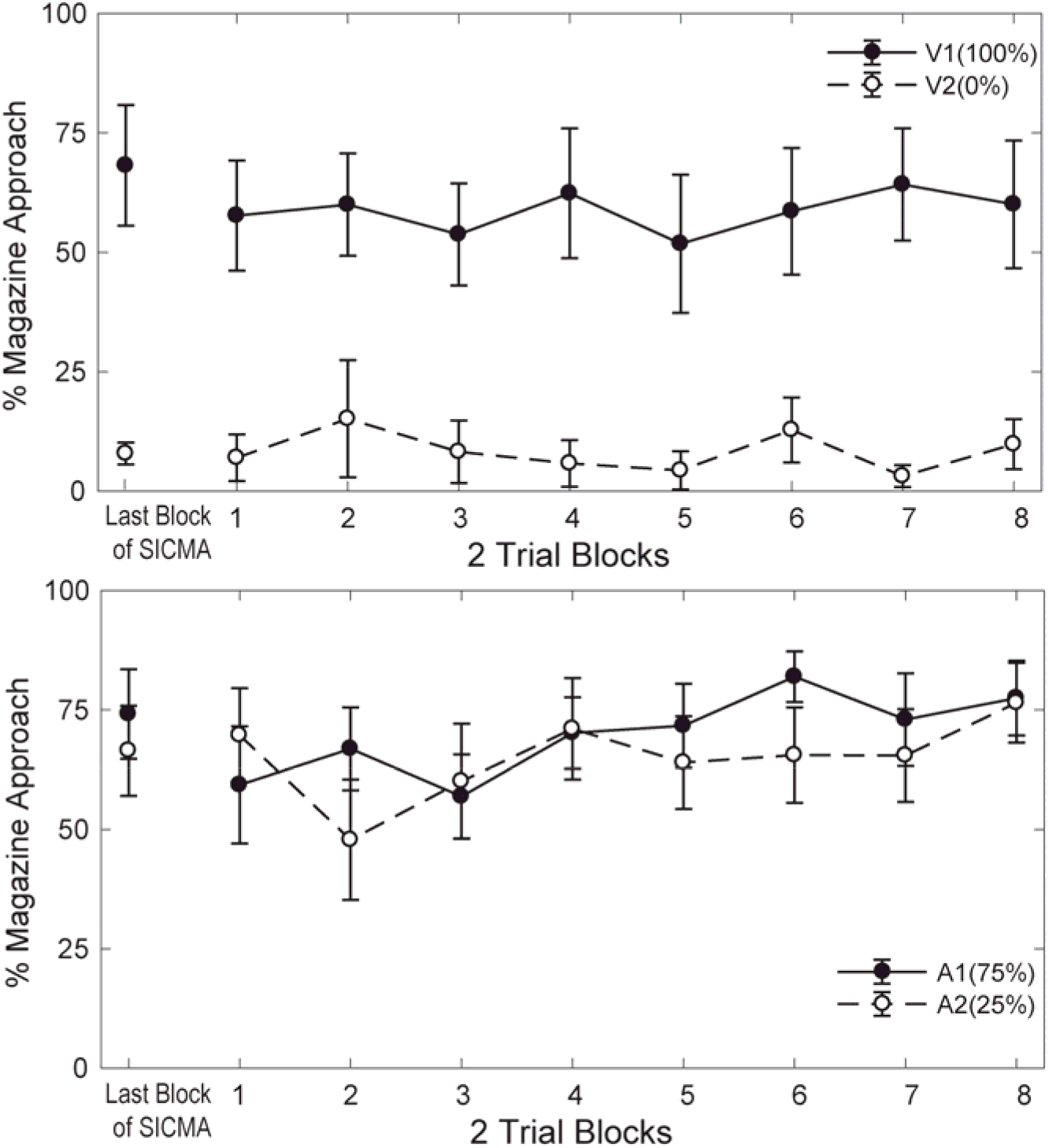
Results of a SICMA-to-Pavlovian transfer test. The same visual and auditory discriminations used in Experiments 1 and 2 were first trained in the SICMA procedure and then tested in a Pavlovian fashion (i.e., without self-initiation). The figure provides a comparison of conditioned magazine-approach performance between the last 2-trial block of the final SICMA session and all 2-trial blocks of the subsequent Pavlovian session. Only data from the last 5 s of cue presentation was considered. Error bars represent the within-subject SEM.

## 5. General discussion

Probing the neural mechanisms of cue-reward learning is often hindered by the difficulty in adapting extant Pavlovian preparations to the parametric requirements of neural recording. In this article, we introduced SICMA, a self-initiated variant of the Pavlovian magazine-approach procedure designed to empower the electrophysiologist working with rodents. Unlike its Pavlovian predecessor, SICMA allows extensive sampling of multiple trial types in a short space of time, leveraging the experimenter’s ability to detect real patterns in the neural data without compromising learning.

A further advantage of SICMA for neural recording is that it guarantees that at the onset of each CS the animal is in the same location within the conditioning chamber. This will help reduce trial-to-trial variability in neuronal responses caused by location-dependent changes in the perception of the stimuli and/or by the juxtaposed encoding of spatial and cue-related information. In addition, SICMA ensures that at the onset of each CS the animal is engaged and thus more likely to consistently garner task-relevant attentional resources that would likely fluctuate across trials over the course of a long Pavlovian session. Indeed, a disadvantage of the standard magazine-approach procedure for neural recording is the possibility that the animal might become oblivious of the CSs as they continue to be presented.

A higher level of engagement in SICMA might go some way to explaining the superior performance observed in this condition relative to the yoked Pavlovian control. However, other explanations should be considered, particularly to account for SICMA’s imperviousness to the detrimental effects of massed trials on learning so typical of Pavlovian conditioning preparations. The latter effects are commonly attributed to lessened extinction of the context due to the high frequency of reinforcement, which will enhance the context’s ability to compete with discrete CSs for behavioral control (e.g., Rescorla, Durlach & Grau, 1985). By making trial initiation contingent upon an instrumental response (e.g. poking in the noseport to turn on the CSs), the role of the context as a predictor of reward might drastically diminish in SICMA. In addition, deleterious memory-interference effects might have less impact on learning in SICMA than in the yoked Pavlovian group. For instance, any proactive interference resulting from lingering short-term memory traces carrying over to the next trial would be attenuated in SICMA if the trial-initiating response can *reset* the short-term memory buffer (Dunnett & Martel, 1990). Alternatively—or additionally—agency over trial-initiation might reduce retroactive interference of each trial with rehearsal of the preceding trial by removing any element of surprise that trial presentation has when delivered in a Pavlovian fashion with a variable ITI. This would place SICMA rats at an advantage over yoked ones in light of evidence that a surprising event presented shortly after a trial can disrupt learning on that trial (Wagner, Rudy & Whitlow, 1973). Future investigations of these mechanisms will not only inform the use of SICMA, but more broadly, shed light on the role of agency in predictive learning.

While the current procedure offers a series of advantages for neural recording, it also comes with some downsides. Notably, the self-initiation aspect of the procedure makes it in principle difficult to apply to the study of aversive conditioning. Even if an aversive component were superimposed on the appetitive task, the number of aversive trials would necessarily have to be relatively small if the animal is not to be discouraged from performing altogether—in all likelihood small enough to represent no advantage over extant aversive procedures. Furthermore, giving the animal control over trial initiation requires a minimum, nonzero overall rate of reinforcement in order to maintain the animal’s motivation to perform. Extensive pilot work in our laboratory has revealed that rats will perform in SICMA for ∼100 trials at a 25% overall reward rate, and it is possible that an even lower rate might support behavior in well-trained animals. That said, it is still the case that SICMA will not be the procedure of choice for studies involving long blocks of nonreinforced trials presented consecutively and with no intervening reinforced trials. Lastly, as hinted above, SICMA will also be of little use to researchers investigating the neural bases of contextual conditioning, as in SICMA the context is rendered unpredictive of reward. Interestingly, eliminating the contribution of contextual conditioning to cue-evoked conditioned responding provides a less ambiguous readout of the cue’s predictive significance (i.e., uncontaminated by context-elicited conditioned responding), which will be advantageous to researchers specifically interested in cue-reward learning.

To the extent SICMA and standard Pavlovian training might engage different cognitive processes (e.g., heightened attention to the task, diminished competition by the context, etc.), one must exert caution when generalizing the results from SICMA studies to Pavlovian settings. The smooth transfer of discriminative performance across the SICMA and Pavlovian phases of Exp. 3, however, tentatively argues for a common discrimination-learning mechanism that informs decision-making under different behavioral requirements. It is upon the neural implementation of that mechanism that SICMA can shed light where Pavlovian preparations fall short. Thus, we anticipate the procedure will be particularly useful in neural recording studies using complex, within-subject discrimination designs (e.g., four trial types or more), such as those typical of stimulus selection, nonlinear discriminations, categorization and rule learning studies.

To conclude, we would argue that a more general limitation of appetitive Pavlovian procedures is that the animal’s role is restricted to that of an opportunistic agent aiming to exploit environmental contingencies beyond its control. By granting the animal agency to seek out cues potentially predictive of reward, the SICMA procedure offers a complementary, also ecologically-relevant way to model appetitive learning.

## CRedit author statement

**Ingrid Reverte:** Conceptualization, Methodology, Software, Investigation, Data curation, Formal analysis, Writing- Original draft preparation. **Stephen Volz:** Methodology, Software, Formal analysis, Visualization, Writing- Original draft preparation. **Fahd H. Alhazmi:** Software, Formal analysis, Visualization, Writing- Original draft preparation. **Mihwa Kang:** Investigation, Validation. **Keith Kaufman:** Investigation, Validation, Software. **Sue Chan:** Software. **Claudia Jou:** Methodology, Software. **Mihaela Iordanova:** Writing- Reviewing and Editing. **Guillem R. Esber:** Conceptualization, Project Administration, Methodology, Supervision, Writing- Original draft preparation, Writing- Reviewing and Editing, Funding acquisition.

## Acknowledgements

This research was supported by National Institute on Drug Abuse, R00 grant 5R00DA036561 (GE) and Brooklyn College’s startup funds (GE). GE would like to thank the Brooklyn College Facilities staff for their help setting up the Esber lab. The authors declare that they do not have any conflicts of interest (financial or otherwise) related to the data presented in this manuscript.

