## Supplemental Materials for "A self-initiated cue-reward learning procedure for neural recording in rodents"

S1. Single-unit recording

To illustrate the importance of sampling cues across multiple trials in neural recording studies of predictive learning, we implanted electrodes in the orbitofrontal cortex of three rats and recorded single-unit activity while the animals performed in a well-trained visual discrimination embedded in the SICMA procedure. After single units were isolated, we conducted a series of bootstrap analyses varying the number of trials sampled, the results of which are shown in Fig. 1.

### S1.1. Materials and Methods

#### *S1.1.1. Animals & Apparatus*

One male (490 grams) and 2 females (227 and 328 grams) Long-Evans rats bred at Brooklyn College from rats of Charles River descent were used. Experimental procedures started when the rats were 6 months old. All husbandry and behavioral apparatus details were identical to those reported in sections 2.1.1 and 2.1.2 of Experiment 1, respectively.

*S1.1.2. Behavioral procedures*

Prior to electrode-implantation surgery, rats received magazine training followed by shaping of the initiating nose-poke response as described in section 2.1.3.1. They then received 10 sessions of discrimination training in the SICMA procedure involving the same visual cues, V1 and V2, used in Experiments 1-3. The discrimination was of the form V1+, V2-, V1V2-, where V1 presentations were reinforced, whereas V2 and compound V1V2 presentations were not. Each of these trial types was presented 25 times. All other details of the SICMA procedure were identical to those described in section 2.1.3.

#### *S1.1.3. Electrode-implantation surgery*

Surgical procedures followed guidelines for aseptic technique. 3D-printed electrode headcaps were previously manufactured in-house, each of which featured two individually-drivable 8-tetrode bundles made of 20-µm-diameter tungsten wires (California Fine Wire, Grover Beach, CA). One date prior to surgical implantation the tetrode bundles were trimmed at a 45° to extend ~1.5-2 mm past the tip of their polyimide housing. They were then immersed in a non-cyanide gold solution (NeuraLynx, Bozeman, MT) and electroplated to a target impedance of ~100 kΩ using the nanoZ^TM^ multi-electrode impedance tester system (White Matter, LLC., Seattle, WA). On the day of surgery, rats were anesthetized using isoflurane and the two tetrode bundles were chronically implanted in both hemispheres dorsal to the orbitofrontal cortex (3 mm anterior to bregma, +/– 3.2 mm laterally, and 4 mm ventral to the brain surface). Immediately before removal from anesthesia rats were given Carprofen (5 mg/kg sq) and triple antibiotic ointment was applied around the incision wound. To control pain, buprenorphine (Buprenex SR, 0.05 mg/kg, s.c.) was injected ~15 minutes after removal from anesthesia. In addition, antibiotic Cephalexin (15mg/kg, p.o.) was administered daily for 15 days post-operatively to prevent infection.

*S1.1.4. In-vivo electrophysiological recording*

After a minimum of 10 days of postoperative recovery time, the rats were retrained for 5 days in the above discrimination to get them accustomed to performing in the SICMA procedure while tethered. Continuous voltage data relative to a ground screw implanted in the rats’ skulls was sampled at 25 kHz using an eCube server acquisition system (White Matter, LLC., Seattle, WA). Signals were amplified and digitized at a headstage mounted within a 3D-printed headcap cemented to the rats’ skull. Neural activity was monitored in real-time from an adjacent room via Open Ephys software (Siegle et al., 2017) to confirm the presence of isolable single units in each rat. If an implant did not show single units, it was advanced 80 µm at the end of the session by turning a screw mounted to a microdrive. Recording began only after tetrodes were driven 4.4 mm ventral to the brain surface after several days of bundle advancement. We recorded neural activity for 10 additional sessions, only the last of which was analyzed for the purpose of the present paper.

*S1.1.5. Spike sorting*

The continuous electrophysiological data were then sorted into isolable single units using the Kilosort algorithm (Pachitariu, Steinmetz, Kadir, Carandini, & D., 2016). Kilosort-produced clusters were then manually curated using the Phy temple-GUI (Rossant, et al., 2016). Spike clusters were labeled as single units if they produced an autocorrelation with evidence of an inter-spike interval, did not show evidence of physical drift in the maximum amplitude of their spikes, and if the amplitude of detected spikes formed at least 90% of a normal distribution (indicating that unit was not submerged in the noise floor). Only clusters that met all of these criteria were considered for further analysis.

*S1.1.6. Histology*

At the conclusion of the experiment all rats were perfused using a phosphate-buffered solution followed by 4% paraformaldehyde (Alfa Aesar, Ward Hill, MA). Brains were then extracted and sliced to 40 μm sections. Sections were then stained with cresyl violet (Alfa Aesar, Ward Hill, MA) for the purpose of verifying electrode placements.

*S1.1.7. Statistical analysis*

All analyses were conducted using the firing rates during cue presentation of 78 neurons drawn from the three rats. This resulted in 75 firing-rate values per neuron, corresponding to 25 presentations of each of the 3 trial types V1+, V2- and V1V2-. We then performed a one-factor ANOVA on each of these neurons to determine which ones significantly discriminated between reinforced and non-reinforced cues. We also calculated the observed statistical power of each neuron that produced a significant F value. We then decimated the number of trials included in the statistical analysis to N=5, 10, 15, and 20, and recalculated the F value with those numbers of trials. Included trials were randomly sampled without replacement. This process was repeated 1000 times per N value.

*S1.2. Results & Discussion*

The top panels of Fig. 1 show the average number of significant neurons (at an alpha value of 0.05, uncorrected) from each rat across bootstrapped iterations as a function of sampled trials. The number of neurons that significantly discriminated reinforced from nonreinforced cues increased as a function of trials. For each additional trial included, the number of neurons that reached significance increased by 2.2%, as demonstrated by regression analysis (p<0.001).

We examined the effect of the number of trials on the statistical power of the ANOVA (Fig. 1, bottom panels). We calculated the observed power, which quantifies the probability of correctly detecting a neuron that discriminates between reinforced and nonreinforced cues. This analysis included only neurons deemed significant at the 0.05 level, using all 25 trials. Then for each of these neurons, we again sampled trials to a decimated total of 5, 10, 15 and 20. For each set of sampled trials and for each neuron, we performed an ANOVA to test the omnibus effect of firing rates across the 3 trial types and then we converted the bootstrapped F-values into observed power using the *powerAOVI* toolbox in Matlab® (Trujillo-Ortiz & Hernandez-Walls, 2002). For each number of sampled trials, the tests were repeated 1000 times with randomly selected samples. Overall, each additional trial increased power by ~3%, as demonstrated by regression analysis (p<0.001).

S2. Supplemental Figures

S2.1. Behavioral apparatus

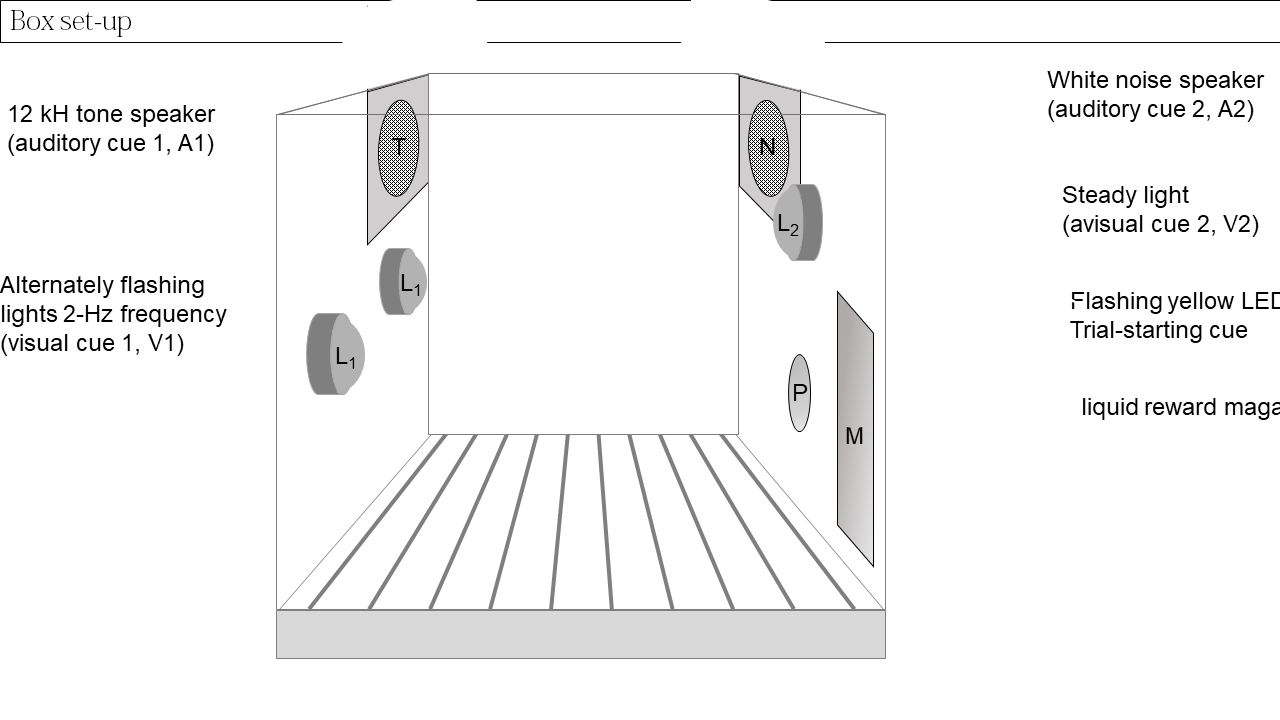

*Figure S1.* Schematic depiction of the interior of the conditioning chambers as viewed through the Perspex front door, specifying the spatial location of all stimuli and response apparatus used. Left wall: T represents a speaker used to play a 12-kHz, 70-dB tone used as one of the auditory cues (either A1 or A2), whereas L1 represents two jewel lights that were flashed alternately at 2 Hz, together forming one of the visual cues (either V1 or V2). Right wall: N depicts a speaker used to play a 70-dB white noise that served as the second auditory stimulus (either A1 or A2), whereas L2 depicts a jewel light used to provide the steady light CS that served as the second visual cue (either V1 or V2). Lastly, P represents the nose-poke port used to perform the self-initiating response, whereas M represents the reward magazine where the sucrose reward could be collected and magazine approach was measured.

S2.2. Rate of head entries as a dependent variable in SICMA-trained rats

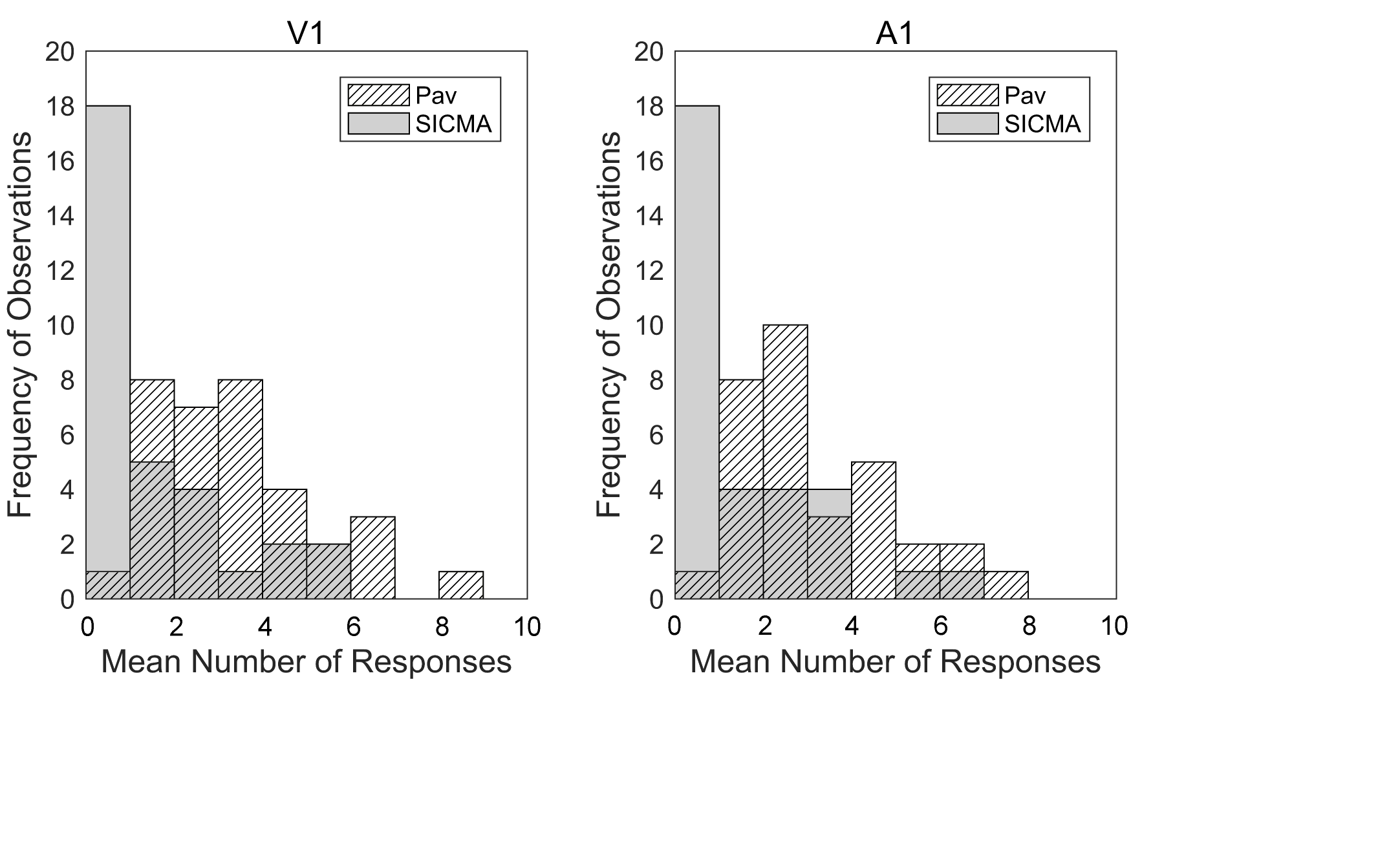

*Figure S2.* Count of rats in the SICMA and Pav groups across the last two sessions of training of Experiment 1 plotted as a function of the average number of head entries in the reward magazine during the 100% reinforced cue V1 (left panel) and 75% reinforced cue A1 (right panel). Both panels suggest that overall SICMA rats made fewer head entries than Pav rats, with most SICMA rats making a single entry during either cue. This difference between the group distributions was statistically significant for both cues.

S3. Supplemental Tables

S3.1. Stimulus counterbalancing

| **Subject** | **A (100%)** | **B (0%)** | **X B (75%)** | **Y (25%)** |
| --- | --- | --- | --- | --- |
| Male 1 | Flashing Light | Steady Light | White Noise | 12-kHz Tone |
| Male 2 | Flashing Light | Steady Light | 12-kHz Tone | White Noise |
| Male 3 | Steady Light | Flashing Light | White Noise | 12-kHz Tone |
| Male 4 | Steady Light | Flashing Light | 12-kHz Tone | White Noise |
| Female 1 | Flashing Light | Steady Light | White Noise | 12-kHz Tone |
| Female 2 | Flashing Light | Steady Light | 12-kHz Tone | White Noise |
| Female 3 | Steady Light | Flashing Light | White Noise | 12-kHz Tone |
| Female 4 | Steady Light | Flashing Light | 12-kHz Tone | White Noise |

Table S1. Stimulus counterbalancing within each squad of 8 rats across all experiments.
